# From optimality to reality: decision-making in basal ganglia output

**DOI:** 10.1101/2025.10.29.685254

**Authors:** Gil Zur, Noga Larry, Mati Joshua

## Abstract

Value-based decision-making requires translating abstract representations of alternative values into concrete decisions. Although this process is widespread, the neural mechanisms underlying this transformation remain unclear. Here, we address this question by examining neural dynamics in the substantia nigra pars reticulata (SNpr) of two monkeys performing a decision-making task. We found a dynamic shift in SNpr activity during the decision process. Before choice, SNpr activity encoded the optimal selection regardless of the eventual action taken, representing the action and value of the optimal choice even when the non-optimal target was ultimately selected. During choice, the SNpr shifted to code the actual selection made by the monkey. This pattern was reinforced by trials with suboptimal choices, revealing a clear transition in SNpr coding from representing the abstract value of the optimal selection to encoding the chosen action, thus suggesting a neural mechanism that transforms value representations into concrete decisions.

## Introduction

Decision-making is a core cognitive function that enables individuals to choose between options based on their expected outcomes. In value-based decisions, maximizing the reward outcome requires maintaining a reward representation to assess the expected reward for each alternative (i.e., action valuation) and selecting the appropriate action to obtain it^1^. While these components of the decision process may be considered separately from a theoretical perspective for convenience, the underlying neural mechanisms are likely more complex. Multiple brain regions may contribute to different subtasks, functioning in parallel rather than as distinct modules with a clear hierarchy. One such potential region is the basal ganglia, which is widely recognized for its role in decision-making^2–5^. On the one hand, the basal ganglia act as a critical reward hub, which together with the frontal cortex, forms a loop involved in reward representation and action valuation^6^. On the other, the basal ganglia are also part of the motor system. Their output structures enable action execution through subcortical circuitry such as the superior colliculus (SC)^7–12^. Indeed, recent studies have reported mixed selectivity in the basal ganglia output structure, the substantia nigra pars reticulata (SNpr), where neurons responded to various task features and exhibited high-dimensional dynamics^13,14^. The SNpr transmits information to both the SC and the cortex—two pathways that may serve distinct functions in the decision-making process. Given its complex temporal dynamics and encoding of both target values and motor commands, the SNpr is a locus of integration in which abstract target selection may develop into a concrete motor command. However, despite its potential, the SNpr has been underexplored in the context of decision-making^9^, especially with regard to the neural signals it conveys during value-based decisions.

In this study, we investigated the role of the SNpr in value-based decision-making. Given the rich temporal dynamics of SNpr activity and its potential role in integrating reward expectation and action encoding, we aimed to explore how these dynamics unfold during the decision-making process. For this purpose, we recorded neuronal activity from the SNpr of two monkeys performing a decision-making task involving eye movements guided by reward probability. We observed dynamic transitions in SNpr coding that shifted from reward representation and the extraction of optimal selections to the coding of actual behavior that did not always align with the optimal choice. We paid particular attention to failure trials, where suboptimal selections were made, since these trials provide a unique opportunity to validate the coding of optimal selection and the subsequent transition to non-optimal behavior.

## Results

To investigate SNpr dynamics during value-based decision-making, we trained two monkeys on smooth-pursuit selection tasks^15,16^ that included reward manipulations, target selection, movement planning, and dynamic movement control (Figure 1A). Each trial began with a white fixation target appearing in the center of a black screen. Next, the cue stage was initiated, during which two colored targets appeared adjacent to the fixation target while the monkey was required to maintain its gaze on the fixation point. The colors of the targets indicated the possible probabilities of receiving a reward upon completing the trial. Each trial involved two different colors chosen from five options, with each color corresponding to a distinct probability (0%, 25%, 50%, 75%, 100%). This resulted in 10 possible combinations, referred to as the *reward condition* of the trial. Within each reward condition, we refer to the target with the higher probability as the *maximal* target and the one with the lower probability as the *minimal* target. After a random cue period, the fixation target disappeared, and the two targets began moving from their eccentric position towards the center of the screen and beyond. The monkey was required to select one of the targets to complete the trial (see below). Upon successful trial completion, the monkey was rewarded according to the probability associated with the selected target. For each reward condition composed of a pair of probabilities, the *direction condition*, which refers to the motion direction of the maximal and minimal targets, was randomly interleaved, resulting in a total of 20 conditions (10 reward conditions × 2 direction conditions).

**Figure 1:**
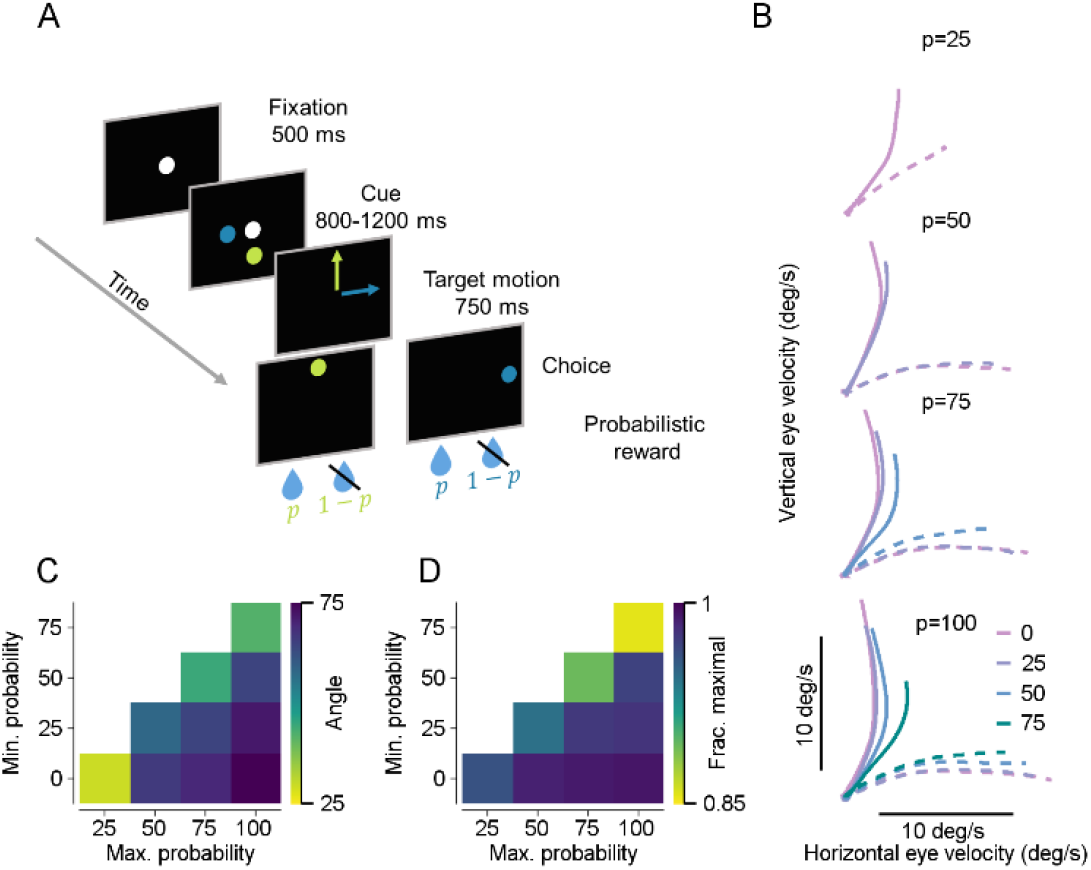
Task and behavior. **A**. Schematics of the tasks. The spots represent the target; the arrows show the direction of motion and *p* corresponds to the probability of receiving a reward. **B**. Averaged eye velocity in the horizontal versus vertical directions in the first 300 ms following motion onset. The value at the top of each panel represents the maximal probability. Colors indicate the minimal probability. Solid and dashed lines depict conditions where the maximal probability target moved vertically and horizontally. **C**. The angular difference between the solid and dashed lines in **B** for each reward condition. The maximal probability is shown along the horizontal axis and the minimal probability along the vertical axis. **D**. Monkeys’ performance in choosing the maximal probability target for each probability pair. Axes are the same as in **C**.

### Behavioral bias and probability-dependent selection during movement execution

In response to the motion onset of the two targets, the monkeys initiated a smooth pursuit movement in an intermediate direction between targets, which was influenced by the reward probability associated with the targets. Figure 1B presents the monkeys’ eye velocity during the first 300 ms of the motion epoch. Plotting the eye’s horizontal against the vertical velocity within this time window reveals a bias of the eye velocity from purely horizontal or vertical tracing toward an intermediate direction^15,17,18^. Typically, this initial biased pursuit was followed by a saccade to the selected target. We detected this saccade online and only kept the selected target, which was then pursued until the end of the trial. Hence, the saccade represented the monkey’s final selection. We refer to the earlier, uncertain phase of pursuit as the pre-commitment stage. This early bias towards the maximal probability target should not be interpreted as the final selection, since even in trials where the minimal probability target was ultimately chosen, the eye could still show an initial orientation toward the higher reward probability (Figure S1). Thus, in this target selection task, information about reward probability not only influenced which target was ultimately chosen but also the dynamics of the motion trajectory.

The magnitude of the pursuit bias increased as the difference between the targets’ probabilities decreased (Figure 1B, see Figure S1 for error trials). To quantify the monkey’s bias for each reward condition, we measured the angle between each pair of matching reward conditions with different directional conditions (Figure 1C). Note that a smaller angle indicated greater bias from the targets, because the monkey was less decisive in selecting a direction. We also quantified the final selection as the fraction of selections of the maximal probability for each reward condition (Figure 1D). The monkeys’ performance in selecting the maximal probability target was above 85% in all conditions, with an average of approximately 95%.

We found two main behavioral effects. First, as the difference between the probabilities increased, the monkeys’ accuracy improved (Figure 1C,D, the colors in each column darken from top to bottom indicating that for each fixed maximal probability performance increased when the minimal probability decreased). Second, for constant probability differences, the size of the probabilities affected the monkeys’ performance (Figure 1D, colors darken from the top-right to the bottom-left corner on the matrix diagonals), as the monkeys were more likely to select the target with the highest probability as the overall probability values decreased. However, this pattern was not observed during the pursuit phase in the pre-commitment stage (Figure 1C).

### Optimal selection encoding in SNpr population activity

We recorded single-neuron activity from the SNpr of two monkeys while they performed the target selection task. Figure 2A depicts a PSTH of an individual SNpr neuron during the cue period, across all reward conditions in both direction conditions. The neuron exhibited responses corresponding to the maximal target within each reward condition, as illustrated by the similarity among curves of the same maximal probability (color-coded). We assessed how strongly individual neurons encoded specific task features during the cue period by calculating the effect size of each neuron^19–21^ (Figure 2B). This analysis decomposed the trial-by-trial variability in neuron activity into components associated with each experimental factor (see Methods). SNpr neurons exhibited a larger effect size for maximal probability during the cue epoch compared to the minimal probability (paired permutation-test, p=0.00002) and direction (paired permutation-test, p=0.0001).

**Figure 2:**
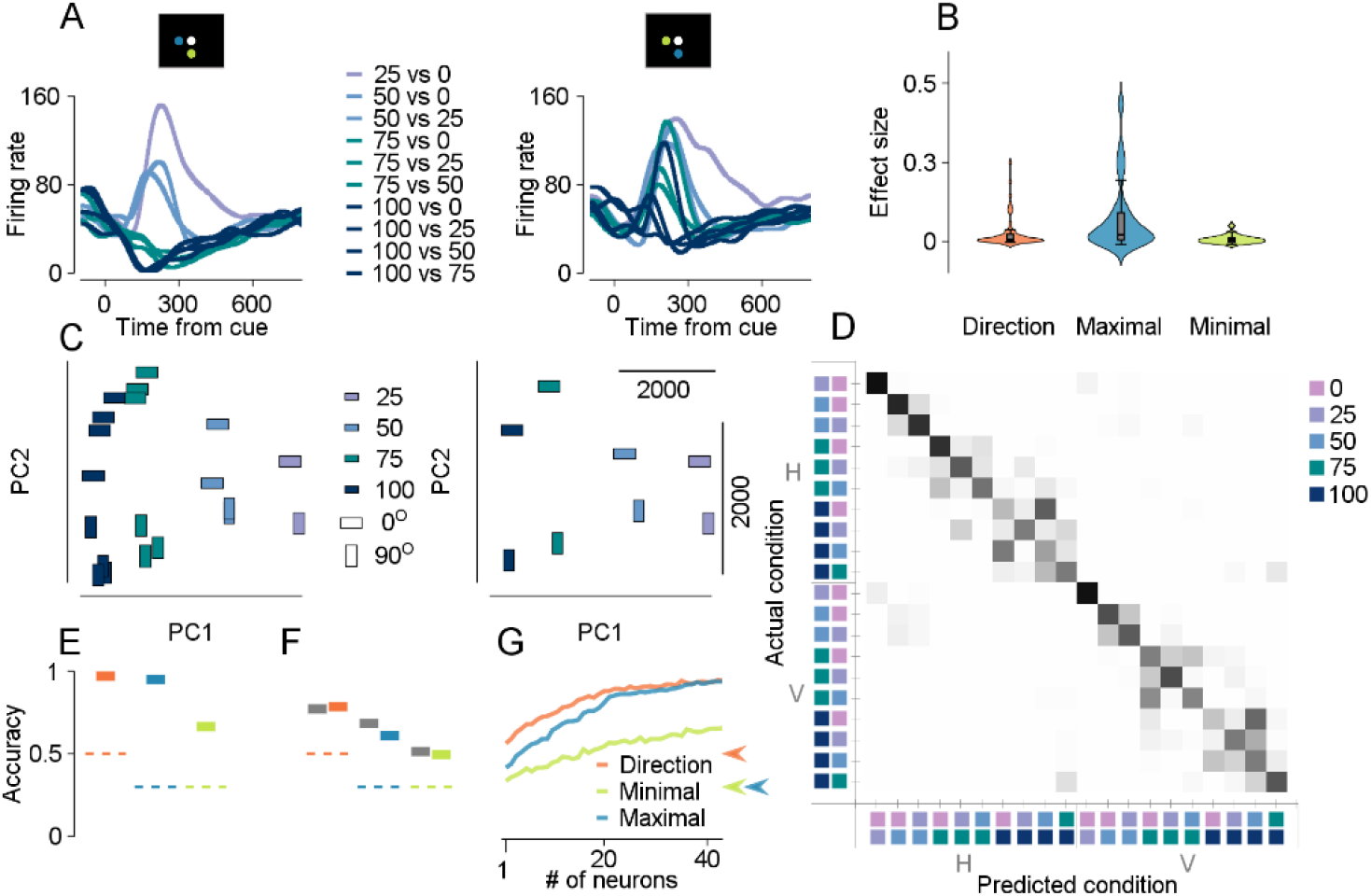
Optimal selection encoding in SNpr population activity. **A**. PSTH illustrates the firing rate of an example neuron during the cue phase. Colors correspond to the maximal probability in each reward condition. The inset screen diagram displays the direction condition (horizontal or vertical), with the blue target representing the maximal probability. **B**. Effect size of SNpr neurons across all experimental factors during the cue stage. **C**. Left: PCA projection onto the first two principal components for SNpr population activity during the cue phase across all task conditions. Color indicates the maximal probability of each condition, and the rectangles’ orientation reflects the direction condition. Right: Same PCA projection as on the left, with each rectangle’s position indicating the mean position of the matched rectangle colors from the left panel. **D**. Confusion matrix displaying the decoder’s prediction accuracy during the cue phase. The intensity of the color indicates the probability of the prediction. The vertical and horizontal axes represent the actual and predicted conditions, respectively. Each axis is divided into two halves, with H and V denoting conditions in which the maximal probability moved along the horizontal and vertical axis. Reward conditions are represented by pairs of colored squares according to the legend, where colors closer to the axis indicate lower probabilities and those further away indicate higher probabilities. The reward conditions are sorted by maximal probability and then by minimal probability. **E**. Decoder accuracy (vertical axis) results for each condition component (horizontal axis) derived from Figure **A**. Dashed lines indicate chance level, and rectangles represent population accuracy. Color indicates the experimental factor (legend in **G). F**. Data-constrained decoder accuracy on error trials for each experimental factor (legend in **G**). Gray rectangles represent the accuracy of the corresponding correct trials serving as a control. Dashed lines indicate chance level. **G**. Decoder accuracy as a function of the number of neurons randomly selected from the entire population. Colors represent different experimental factors, with arrows indicating the corresponding chance level for each factor.

To further explore the information represented in the major components of SNpr activity, we performed principal component analysis (PCA) on the population of SNpr neurons (Figure 2C). The first PC (PC1), accounting for 41% of the total variance, primarily captured the maximal probability of the reward condition (color-coded along the horizontal axis), whereas the second PC (PC2), explaining 17% of the variance, primarily reflected the direction condition (represented by the shape angle along the vertical axis).

Because effect size is indicative of the single-neuron coding and the PCA is indicative of the main components of population activity, both analyses provide a reduced data representation. To gain a fuller understanding of population neural activity, we designed a classifier (see Methods) and applied it to the cue epoch of the decision-making trial. To reveal the coding schemes of the SNpr neurons we generated pseudo-population activity by selecting a single trial from each neuron and combining trials across all neurons. We then decoded the trial condition from the full population activity (see Methods). We randomly repeated this process numerous times for all task conditions and calculated the confusion matrix of the decoder. Figure 2D presents the confusion matrix of the population decoder during the cue stage. The structure of the confusion matrix indicates which information is coded by the neurons. Notably, the decoder demonstrated highly accurate coding of the direction condition, as shown by the blank squares off the main diagonal, which occur where the predicted and actual direction conditions differ. To further evaluate the coding strength and population size required to decode each condition component, we derived the decoder accuracy for each experimental factor using the confusion matrix (Figure 2E). This was done by summing, for each row in the confusion matrix, the entries where the actual and predicted values of the variable matched, and subsequently averaging these row-wise sums. We found that direction and maximal probability were nearly perfectly decoded from the SNpr population, even with a small subset of neurons (Figure 2G), while decoding of maximal probability was more accurate compared to minimal probability. This underscores the SNpr’s preferential coding of the optimal selection option.

So far, we have only analyzed trials in which the monkey selected the target with the maximal probability (*correct* trials). Next, we extended our analysis to trials in which the monkeys’ final selection was sub-optimal (i.e., the minimal-probability target was chosen), which we refer to as *error* trials. Our goal was to determine whether SNpr activity in these trials resembled that of trials where the monkeys selected the optimal target or whether it aligned with the eventual erroneous choice. We developed a decoder (see Data-Constrained Decoder in Methods) trained on the correct trials and tested it on the error trials. Due to the relative sparseness of the error trials, not all neurons had data for each condition in these trials. Therefore, the number of neurons available for training the decoder was constrained by the availability of error trials, resulting in a decoder with less statistical power. To account for this, we also tested the decoder on a subset of correct trials as a control to validate its performance on the error trials. The performance of this decoder in reading out task components was similar on the error and correct trials (Figure 2F). The decoding of the maximal target position was comparable, and representation of the maximal probability remained stronger than that of the minimal probability across both trial types. These findings suggest that SNpr neurons consistently emphasized the maximal probability, even when the monkey ultimately chose the target associated with the minimal probability. Thus, the SNpr appeared to extract the optimal selection for each trial, independent of the monkey’s final choice.

### SNpr activity during motion explains the effect of reward on behavior and final selection

The SNpr’s response during the cue period corresponded to the optimal selection, even on trials where the subsequent choice was non-optimal. This finding is particularly notable, as it suggests an abstract representation of reward that is independent of the actual selection, despite the assumed basal ganglia’s assumed role in action selection and the expectation that SNpr activity would align with the chosen action. We thus next examined SNpr activity during the action period. Unlike the cue period where SNpr activity was primarily driven by the maximal probability, the motion phase showed a significant shift, with direction coding becoming more dominant over the maximal (paired permutation-test, p=0.004) and minimal (paired permutation-test, p=0.00002) probabilities, as reflected in the effect sizes of individual neurons (Figure 3A). Examining the PCA of the population during the motion stage revealed a reversal of the pattern observed in the cue period (Figure 3B): PC1 primarily captured the direction condition (41% of the variance), while PC2 accounted for reward probability (14% of the variance).

**Figure 3:**
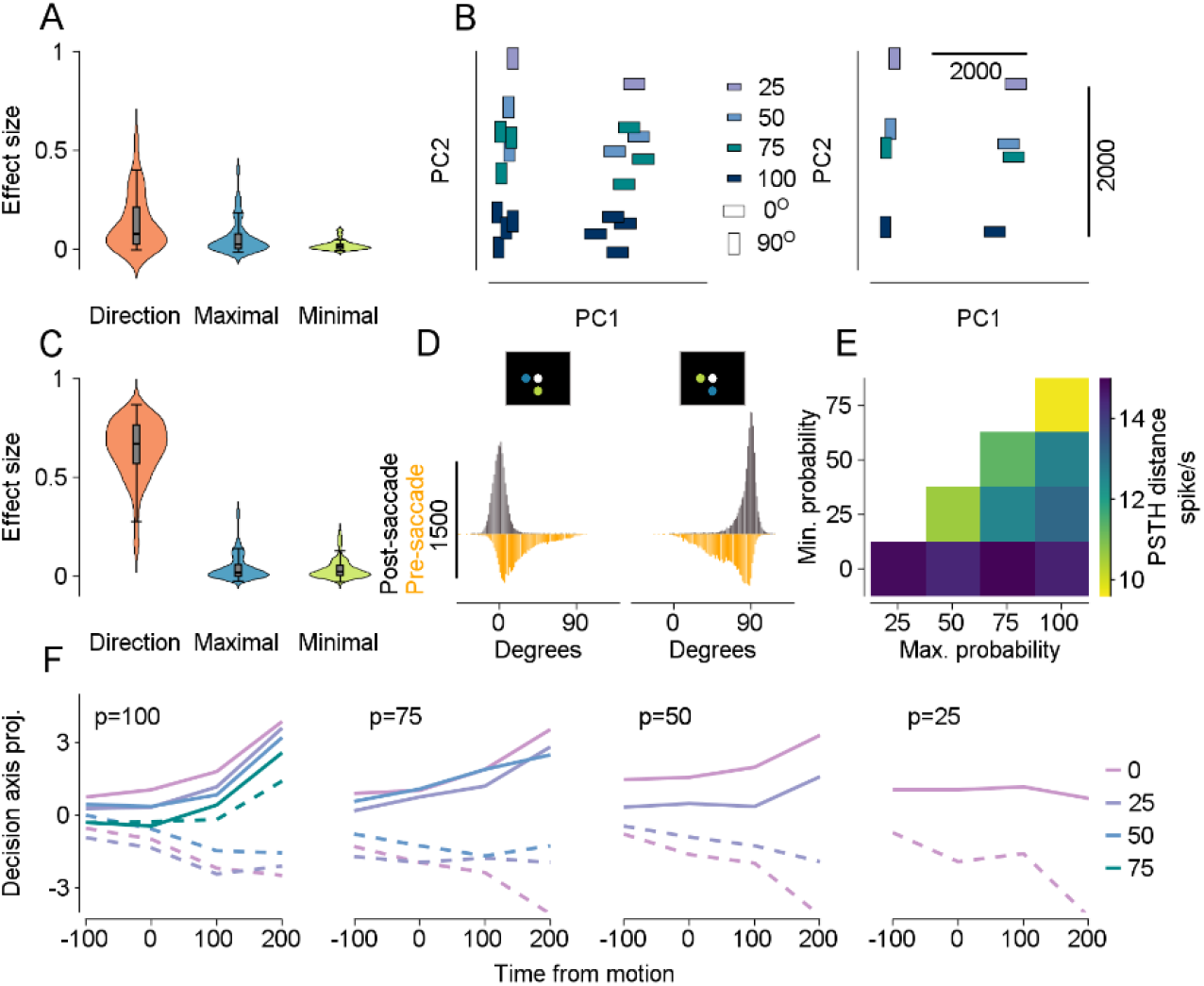
SNpr activity during motion explains the effect of reward on behavior and final selection. **A**. Effect size of SNpr neurons across all experimental factors during the motion stage. **B**. Left: PCA projection onto the first two principal components of SNpr population activity during the motion phase across all task conditions. Color indicates the maximal probability of each condition, and the rectangle angle reflects the direction condition. Right: Same PCA projection as on the left, with each rectangle’s position indicating the mean position of the matched rectangle colors from the left panel. **C**. Effect size on eye trajectory angle for each experimental factor. **D**. Histograms of eye velocity angle (degrees) calculated over 200 ms before (orange) and after (gray) the saccade. **E**. Euclidean distance between PSTHs for direction conditions within each reward condition, measured within a time window of −300 to −100 ms relative to the commitment saccade. The maximal and minimal probabilities are shown on the horizontal and vertical axes. Units are in spikes per second (see Methods, PSTH distance). **F**. Projection of SNpr population activity onto an axis defined by the coefficient of the direction regressor of the single neurons. The horizontal axis represents time from target motion onset. The value at the top of each panel represents the maximal probability, while colors indicate the minimal probability. Solid and dashed lines depict conditions where the maximal probability target moved vertically and horizontally.

The above results reflect SNpr coding across the entire motion epoch. However, during the initial motion period, the eye trajectory differed from the later stage. Examining eye movement angles before and after the saccade on correct trials revealed that while the movement direction approached the selected target before the saccade, full alignment only occurred afterward (Figure 3D; see error trials for comparison in Figure S8). We observed that all the experimental factors—direction, maximal probability, and minimal probability—influenced initial eye trajectory and selection (Figure 1B-D). Therefore, we took a closer look at the pre-commitment stage of motion. To quantify each factor’s impact on behavior, we measured the effect size on the motion trajectory angle during the final 50 ms of the trajectories, as shown in Figure 1C. We found comparable effect sizes between the two reward probabilities (paired permutation-test, p=0.27), whereas the directional experimental factor had a much larger impact (Figure 3C, paired permutation-test: direction vs. minimal/maximal, p=0.00002). Thus, the pursuit trajectory at the pre-commitment stage was guided here by both targets.

We found that the impact of direction was dominant during the motion period in terms of both behavior (Figure 3C) and neural activity (Figure 3A). Therefore, we expected to observe a closer correspondence between neural activity metrics and behavior. Specifically, we expected that conditions that were far apart in terms of behavior would also be more different in terms of neural activity. To test this, we examined SNpr activity just before the saccade, aligned to the commitment saccade time window (−300 ms to −100 ms before the saccade), to identify corresponding neural evidence linked to the monkeys’ selection. For each pair of conditions with the same reward condition but with different direction conditions, we computed the distance between the PSTHs in this pre-saccadic window (see Methods), resulting in a distance matrix for each probability pair (Figure 3E). This analysis revealed two patterns that mirrored the behavioral findings. First, within each maximal probability, the PSTH distance decreased as the difference between probabilities decreased, reflecting the behavioral effect where a larger difference between probabilities led to more accurate target selection. Second, for a fixed difference between probabilities, smaller probabilities were associated with larger PSTH distances, reflecting better separability between the conditions and resulting in more accurate selections by the monkeys (as seen in the diagonals of Figures 3E and 1D). These results were not observed when performing the same analysis during the cue phase and were only partially observed in the early stage of the motion phase (Figure S3). Together, these neural results align with the observed behavioral patterns in the monkeys’ final selection, supporting the hypothesis that SNpr activity plays a key role in driving behavior in value-based decisions.

In the PSTH difference analyses described above, we used each cell’s full temporal pattern of activity. This allowed us to reveal coding patterns while minimizing the assumption regarding the activity pattern containing the information. We next conducted a complementary analysis aimed at identifying population patterns along specific axes of interest^22,23^ to examine the dynamics of the SNpr during the pre-commitment stage more closely. Each neuron was fitted with a linear model (see Methods), allowing us to project SNpr population activity onto an axis defined by the vector of the direction condition coefficients across all neurons (see Figure S4 for cue epoch). We coupled this analysis with a leave-one-out procedure to mitigate potential statistical biases. We found that the dynamics of the SNpr aligned with the monkeys’ behavior (Figures 3F and 1B): as the final selection moment approached, the activity trajectories corresponding to the same reward condition across different directions began to diverge more markedly as the difference between the probabilities increased. Thus, SNpr activity reflected the influence of reward probability on eye trajectory during the pre-commitment stage. Furthermore, population activity projected on a single axis resembled behavior, suggesting that simple linear readouts of the SNpr activity may drive behavior (Figure S5).

### SNpr activity transitions from representing the optimal selection to encoding the actual selection

We found that SNpr activity predominantly reflected the optimal selection during the cue phase while aligning more closely with behavior during movement. Therefore, to examine the time course more precisely, we next examined the dynamics of SNpr activity at each task stage by tracking our decoder’s performance over time (Figure 4A,B). For direction, the SNpr exhibited strong coding throughout all task stages (Figure 4A). However, for reward probabilities, the SNpr displayed a more intricate dynamic pattern. In particular, during the cue stage, the maximal probability was encoded much more strongly than the minimal probability, and achieved nearly perfect coding (Figure 4B). This trend altered during the motion stage. In the pre-commitment phase, maximal probability coding decreased compared to the cue stage, while minimal probability coding increased. This shift was also evident when comparing the effect size at cue presentation and motion onset for each target (paired permutation tests: maximal probability, p = 0.0008; minimal probability, p = 0.006; Figure S2). As a result, both probabilities were encoded more similarly just before the commitment saccade (see the histogram in Figure 4B indicating saccade timing), akin to their comparable influence on behavior (Figures 1B, 3C). After commitment to the maximal probability target, coding of the non-selected (minimal) probability decreased, while coding of the selected (maximal) probability remained robust. This trend became even more pronounced when aligning the decoder to the commitment saccade (Figure 4B. right). Thus, SNpr dynamics shifted from encoding the maximal reward and its direction during the cue to aligning with actual selection during movement, mirroring the impact of reward on behavior.

**Figure 4:**
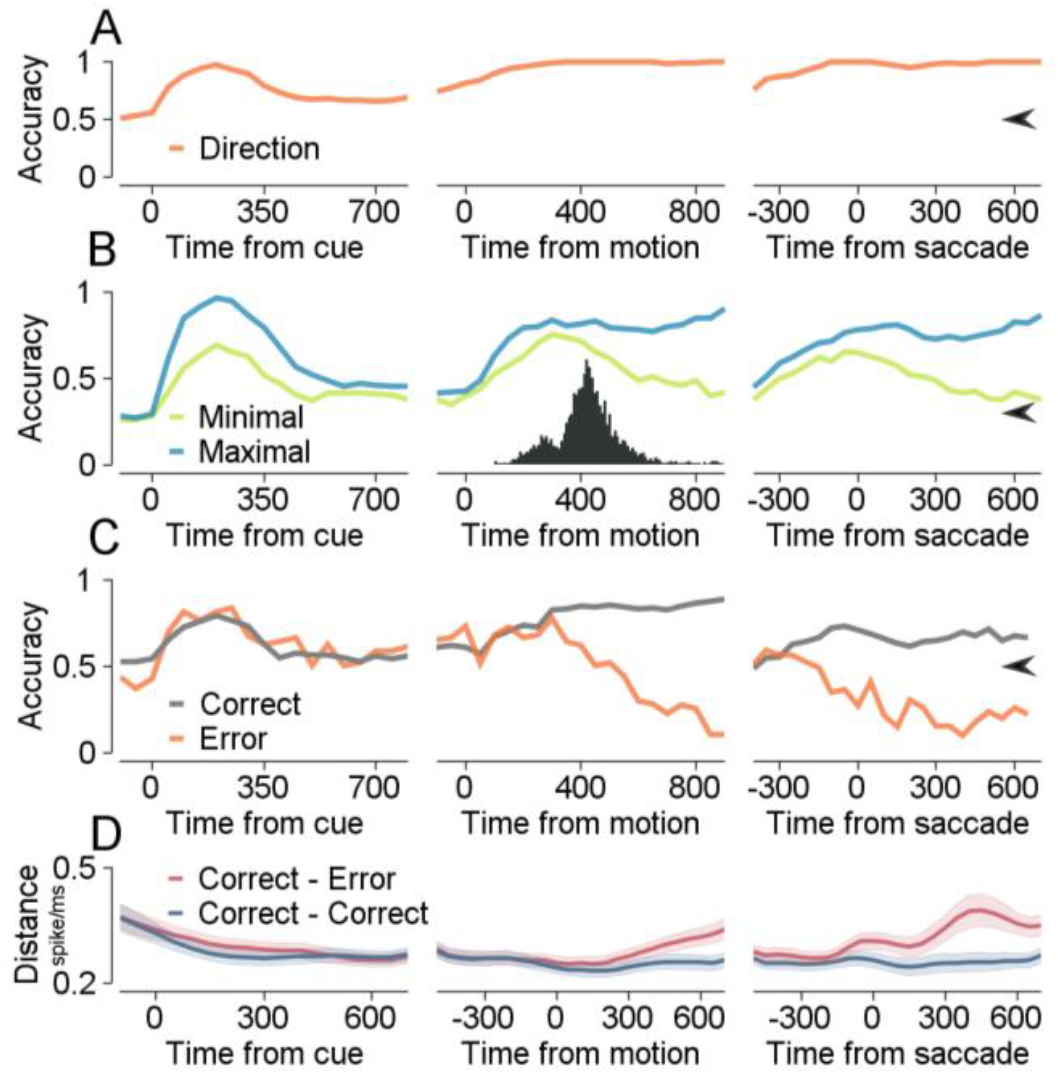
SNpr activity transitions from representing the optimal selection to encoding the actual selection. **A**. Population decoder accuracy for direction condition over time, aligned to cue onset (left), motion onset (middle), and saccade (right). The arrow indicates chance levels. **B**. Same as A, but for reward probabilities. Histograms indicate saccade timing. Colors represent different experimental factors. The arrow indicates chance levels. **C**. Data-constrained decoder accuracy for direction condition across different task stages. Orange and gray lines represent testing on error and correct trials, respectively. The arrow indicates chance levels. **D**. Euclidean distance (vertical axis) over time for different trial stages. Colors represent the groups of trials on which distances were computed.

SNpr activity during error trials at the cue stage initially resembled the optimal selection (Figures 2 and 3) diverging from the eventual behavior. However, due to the strong correlation between SNpr activity and behavior observed during the motion phase (Figure 4), we hypothesized that SNpr coding would shift on the error trials. Specifically, we anticipated transitioning from representing the optimal choice early on to coding for the sub-optimal target as the final selection was made. To test our hypothesis as to a transition in SNpr coding, we applied our data-constrained decoder to both the correct and error trials during the task stages (see Methods). If the hypothesis is correct, during the motion stage, the decoder would classify the selected direction correctly on the correct trials (as seen in Figure 4A) and consistently misclassify the direction on the error trials. As illustrated in Figure 4C, the decoder initially processed both correct and error trials similarly during the cue and the initial phase of the motion stage. However, as time progressed, the decoder accurately classified the direction on the correct trials, while the classification for error trials dropped. Importantly, the classification performance fell below the chance level on the error trials, indicating a similarity in activity between error trials and correct trials in which the monkey moved in the same direction. This further confirms that during movement, the SNpr transitioned to encoding the selected direction rather than the optimal selection on both trial types.

The decoder performance for error and correct trials during initial motion stage was similar, despite the behavioral differences (compare Figures 3D and S8). To better identify when SNpr activity transitioned from coding the optimal choice to the actual choice, we performed a PSTH distance analysis (calculated as the Euclidean distance of the activity pattern, using a bin size of 200 and a step size of 50, see Methods). For each correct trial, we matched either correct trials (Correct-Correct) or error trials (Correct-Error) from the same experimental condition. The Correct-Correct distances served as the baseline that reflected neural activity differences when both the experimental conditions and the monkey’s choice were identical. In contrast, the distances in the Correct-Error case could arise from the same sources as in the Correct-Correct case or from differences associated with the final selection. Comparing these distances provided direct measures as to when SNpr started to code error and correct trials differently.

We analyzed the similarity in SNpr activity across different time epochs to determine how neural patterns diverged between Correct-Correct and Correct-Error. The results revealed similar SNpr activity during the cue stage (Figure 4D). As the motion stage began, the coding for error and correct trials started to diverge. Note that aligning the data to the motion onset introduced variability in the timing of saccades across trials, which could contribute to the gradual divergence between the groups. When aligning the comparison to the saccade (Figure 4D, right), the coding of error trials diverged markedly from correct trials just before the saccade. Thus, as the motion phase began, the SNpr transitioned from coding correct and error trials similarly to coding them differently, with the two types of trials differing solely in their final selection behavior. Therefore, this moment may represent the critical point at which the SNpr initiated a decision in error trials, suggesting a potential neural marker for decision-making failures where behavior diverged from the optimal selection. Since we observed coding of the optimal target during the cue stage for both trial types, these results emphasize the SNpr’s shift from encoding the optimal target to encoding the actual behavior of the monkeys.

## Discussion

The findings showed that neural activity in the SNpr reflected the optimal decision early in the trial, even on trials where a sub-optimal decision was ultimately made. Later in the trial, during the movement epoch, the SNpr shifted to encoding the selected target regardless of its optimality. One major issue when interpreting the decision process is the level of abstraction of the representations. For example, activity could be linked to visuospatial and motor properties, such as the location of the stimuli or the direction of movement. However, a more abstract representation that can more easily be generalized, should represent value-based parameters independently of specific sensorimotor features. Our finding that SNpr activity during the cue was primarily related to the maximal reward probability target, rather than the direction of the stimuli or lower probability options (Fig. 2) is indicative of this level of abstraction. This type of representation is found for example in high-order areas such as the orbitofrontal cortex of monkeys^24^, but interestingly not in mice^25^.

Considering its activity during the cue epoch might suggest that the SNpr is part of the valuation system rather than driving behavior directly. However, the findings observed during the movement epoch challenge this view. In the movement epoch, the coding of direction, which is the major contributor to behavior, increased (Fig. 3) and became more dominant than the reward probability (Fig. 2 and Fig. S2). Furthermore, movement details could be extracted from SNpr activity (Fig. 3). Thus, the SNpr shifted dynamically from a more abstract representation of the reward value at the cue to a behavior-based code during movement.

The dual role of the SNpr in representing reward abstraction and coding behavior may reflect its involvement in decision-making within the basal ganglia-cortex loop, as well as its role in motor control^9,26^. However, while mixed selectivity can provide a computational advantage on cognitive tasks^27,28^, it may also disrupt the ability of downstream regions to extract relevant information accurately. One potential solution could lie in the orthogonal representation of different types of information. Previous studies have reported orthogonal representations of sensory input and memory as a potential mechanism to minimize interference between representations^29^. We thus conducted a complementary analysis to assess the separability of reward and behavioral information and found that these aspects were represented in orthogonal directions (see Methods and Figure S9). Thus, despite the complex mixed selectivity of the SNpr, relevant task-related information could still be read out linearly without disruption from the non-relevant components.

Selection errors in our task allowed us to look for the neural correlates of target selection. We identified the first neural signature of errors in the SNpr during the motion epoch. The early similarity of SNpr activity between the correct and error trials during the cue imposes strong constraints on selection in error trials and suggests two options. One possibility is that selection occur during the pre-commitment stage, not during cue presentation. This option implies that during cue presentation, while a full evaluation of the reward expectation is available in the SNpr, including extraction of the optimal selection, the monkeys had not yet decided which target to pursue in the subsequent movement stage.

Alternatively, selection occur at an earlier stage of the task, but evolve outside the basal ganglia or in other pathways within the basal ganglia. For example, activity in areas of the frontal cortex alternates between states associated with the value of two available options^30^. These alternations may result in errors. The possible alternative suggests that selection in error trials within the basal ganglia could mainly occur at a late stage, just before saccade execution. However, given that the SNpr encodes the optimal selection during the cue phase of the task, the source of the failures in some trials remains puzzling. One potential contributing factor could be momentary distraction at movement initiation, which could lead to suboptimal behavior that impacted the subsequent decision. Our analysis revealed that in approximately 22% of the error trials—compared to only 2% of the correct trials—a micro-saccade was initiated very early in the pre-commitment phase (within 250 ms from motion onset). This suggests that early performance deficits may have contributed to selection failures. However, these premature saccades do not account for the majority of the observed error trials.

A complementary explanation is that internal processing distracts the monkeys from correctly evaluating the reward probability at motion onset. As the movement phase begins, the SNpr reallocates computational resources, prioritizing movement-related encoding over the maintenance of probability representations (Figure 2). Neural activity during the pre-commitment phase suggests a transitional period in which resources shift from reward estimation during the cue phase to action encoding in the movement phase (Figures S7). This pattern implies that errors may arise from a competition in resource allocation, wherein the neural processes required for motion encoding interfere with the previously well-established probability representation. Consequently, despite the initial encoding of the optimal selection, the disruption introduced by movement-related processing may lead to errors in some trials. Further research is needed to determine the extent to which these disruptions in selection arise from attentional lapses, resource allocation constraints, or broader mechanisms governing action selection in the basal ganglia.

## Methods

All experimental procedures were approved by the Institutional Animal Care and Use Committees of the Hebrew University of Jerusalem. Data were collected from one female (Monkey G) and one male (Monkey A) Macaca Fascicularis monkey that had been prepared for behavior and neural recordings using techniques described in detail elsewhere.^31^ Briefly, we implanted head holders to restrain the monkeys’ heads and trained the monkeys to track spots of light that moved across a video monitor positioned in front of them (55 cm and 63 cm from the eyes of monkeys A and G). We used liquid food rewards (baby food mixed with water and infant formula) from a tube placed in front of them to reward the monkeys for accurate tracking of the targets.

The position of the eye was continuously tracked using a high temporal resolution camera (Eyelink 1000 - SR research) at a frequency of 1 kHz and the data were collected for further analysis. To record from the basal ganglia, we performed surgery to place a 19 mm diameter cylindrical recording chamber over the brain regions. Based on MRI imaging, we estimated the location of the SNpr in the recording chamber. Then, we lowered the electrodes to this location. At the targeted site, we identified neurons with a high baseline firing rate and the typical extracellular shape of SNpr neurons^32^. We confirmed that some of these neurons exhibited a clear pause during saccades in certain directions. On some recording days, we also identified neurons with pronounced eye position sensitivity, as expected from the neighboring third oculomotor nerve. We used the Mini-Matrix System (Thomas Recording GmbH) to lower quartz-insulated tungsten electrodes (impedance of 1–2 Mohm) into the SNpr. The neural signals were digitized at a sampling rate of 40 kHz using the OmniPlex system (Plexon) and sorted offline (Plexon spike sorter). We only used neurons that displayed distinct clusters of waveforms in the sorting procedure. The sorted spikes were converted into time stamps with a resolution of 1 ms and were inspected again visually to identify any instability or obvious errors in the sorting procedure. To control for potential behavioral differences between reward conditions that might affect our results, we also recorded the monkeys’ licking behavior. We used an infrared beam to track the licks. Monkey A tended not to extend its tongue, so we recorded its lip movements instead.

### Experimental design

#### Task

Each trial began with a fixation stage, where a bright white target appeared in the center of the screen for 500 ms. During this stage, the monkey was required to maintain fixation on the target to initiate the trial (Figure 1A). Following the fixation stage, the cue stage began. Two different colored targets, chosen from a set of five options (purple, green, red, yellow, and blue), appeared 4° from the fixation target. These targets remained on the screen for a randomized duration of 800–1200 ms, which varied between trials. During this stage, the monkey was required to maintain fixation on the central target. The color of each cue signaled the probability of receiving a reward (0, 0.25, 0.5, 0.75, or 1) if the monkey tracked that target in the subsequent motion stage. The probability-color associations were randomized between the two monkeys. At the end of the cue stage, the fixation target disappeared, and the two colored targets began moving at a speed of 20°/s. The targets moved for 750 ms starting from their initial positions toward the center of the screen and continued beyond it. Typically monkey initiate a smooth pursuit movement in an intermediate direction between targets and then saccade to one of the targets. Online we detected this saccade as an eye velocity that exceeded 80 deg/s. After the saccade, only the target closest to the eye remained and the monkey was required to pursuit it to receive a reward. In cases where the monkey did not make a saccade during target motion, at the end of the motion epoch only the target closer to the eye remained. Upon successful trial completion, the monkey was rewarded based on the probability associated with the selected target’s color. After the motion phase ended, the target remained stationary for an additional 500–700 ms. If the monkey’s gaze remained within a 3–5° x 3–5° window around the target, it received the reward as specified by the target’s color. Monkeys were trained on the task for several months before we recorded neural data.

### Data Analysis

The analyses were conducted using Python and MATLAB. To ensure robust results, only neurons recorded for a minimum of 160 trials were included. This criterion resulted in a total of 52 neurons available for analysis, consisting of 25 neurons from monkey A and 27 neurons from monkey G.

For analyses requiring calculations around the selection saccade time points, trials without a selection saccade were excluded. If a neuron had at least one condition where all trials lacked a saccade, the neuron was excluded from these saccade-related analyses. This criterion led to the removal of 9 neurons from the saccade-based analyses.

#### PSTH calculation and distance

To compute the Peri-Stimulus Time Histogram (PSTH) for a neuron, the trials were segmented into the predefined time window. Each trial was then smoothed using a Gaussian filter with a standard deviation of 30. The root mean square (RMS) of the difference between the PSTHs was then calculated to quantify the difference between average responses as follows:

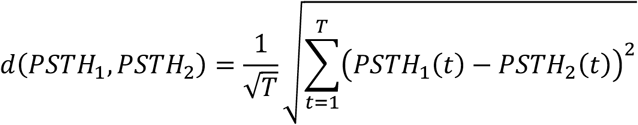

where *N* is the number of neurons, *T* is the duration of the time window in ms resolution, and *PSTHi*(*t*) is the value at time point *t*. This calculation yielded the delta activity between the responses in units of spikes per second. Dividing by 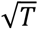 ensured that the result represented the average neural difference in activity across the time window.

To assess the PSTH activity difference at the population level, the PSTHs for all neurons were concatenated before calculating the RMS (see Figures 3, S3).

#### Effect Size

To calculate the coding of a variable throughout the entire epoch, we calculated the partial omega square 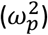 effect size. The partial effect size measures how much of the variability of a neuron is related to the experimental variable in comparison to the variability that cannot be explained by any of the variables. Here we used 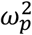 to account for the magnitude of each SNpr neuron’s response to individual aspects of the task conditions, such as target direction, maximal probability, and minimal probability. Specifically, we calculated spike counts in 200 ms bins during the 0–800 ms window following cue (Figure 2B) or motion (Figure 3A) onset. For the pre-saccade period (Figure S2), we analyzed the first 400 ms window after motion onset, excluding trials with early saccades from the analysis. We fit an ANOVA model that included the direction of target motion as a variable, with the addition of time (the specific bin the sample came from). We used the following formula:

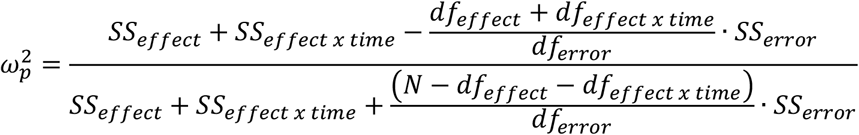

where *SS*_*effect*_ is the ANOVA sum of squares for the effect of a specific variable, *SS*_*error*_ is the sum of squares of the errors after accounting for all experimental variables, *df*_*effect*_ and *df*_*error*_ are the degrees of freedom for the variable and the error respectively and *N* is the number of observations (number of trials x number of time bins). *SS*_*effect x time*_ is the ANOVA type II sum of squares for the interaction of a specific variable (i.e., target direction) with time, and *df*_*effect x time*_ are the corresponding degrees of freedom. We included the interaction term since it quantifies the time-varying coding of the variable.

#### PCA

To perform principal component analysis (PCA) on the SNpr population, we first calculated the PSTH for each individual neuron during the cue (0–800 ms) and motion (0-750) epochs. Next, for each condition, we concatenated the PSTH responses of all neurons, resulting in an *M*_*c*×*nt*_ matrix, where *c* is the number of conditions (20), n is the number of neurons and *t* is the epoch length (e.g., 800 ms for the cue epoch). PCA was then applied to this matrix, and the scores for each condition were plotted on the first two principal components, PC1 and PC2 (Figure 2C).

#### Decoder

To better understand the coding properties of the SNpr, we designed a decoder and applied it to different task stages. For each iteration, we computed the PSTH for each neuron across all 20 conditions (*c*) within a specified time window of length *t*. We then applied a leave-one-out (LOO) approach, randomly removing the activity of a single trial for each condition. This resulted in two matrices for each neuron:

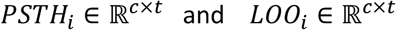

where *i* is the index of the neuron.

Next, we concatenated the PSTH and LOO matrices of all neurons along the column dimension to obtain two matrices representing the population activity:

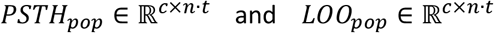

where *n* is the number of neurons, and the resulting dimensions are determined by the number of conditions (*c*) and the product of the time window length (*t*) and the number of neurons (*n*).

To evaluate the decoder’s predictions, we computed the Euclidean distance for each condition (row) in *LOO*_*pop*_ relative to all rows in *PSTH*_*pop*_. The Euclidean distance between the *j*-th row in *LOO*_*pop*_ and the *k*-th row in *PSTH*_*pop*_ is given by:

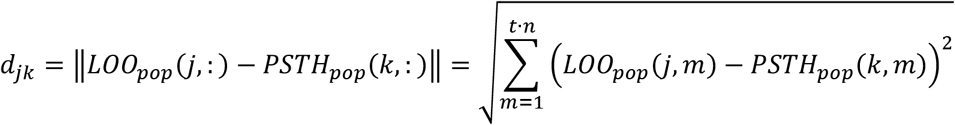

The decoder assigns the predicted condition label for the *j*-th row in *LOO*_*pop*_ as the index of the row in *PSTH*_*pop*_ that minimizes the distance:

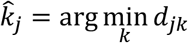

To ensure robustness, we repeated this procedure over 100 iterations. In each iteration, the leave-one-out step was performed randomly, resulting in multiple predictions for each expected condition (true label) corresponding to the number of iterations. Using the multiple predictions for each condition, we constructed a confusion matrix (Figure 2D). The confusion matrix represents the probability of predicting each condition for a given expected condition. Each row in the matrix corresponds to a true condition, while each column represents a predicted condition. The values in the matrix are calculated as the proportion of times a condition was predicted for a given true condition, normalized over all iterations.

#### Data-Constrained Decoder

The procedure for this decoder was similar to the decoder described earlier, with modifications to the test groups and the number of neurons utilized in the creation of the *PSTH*_*pop*_ matrix. Specifically, the control decoder was trained on trials where the monkey correctly selected the maximal-probability target (Correct trials) and tested on trials where the minimal-probability target was selected (Error trials). Given the relative sparsity of error trials, not all neurons contributed data for every condition in these trials, leading to constraints on the training data and a reduction in the decoder’s statistical power. To account for this limitation, we also tested the decoder on a subset of correct trials to serve as a control group for performance on the error trials.

For a given condition *c*, we identified the subset of neurons *N*_*c*_ that had at least one error trial in that condition. Only these neurons were included in the PSTH population matrix for that condition. For each neuron *n* ∈ *N*_*c*_, the number of error trials available for condition *c* was small in comparison to the correct trials. This allowed us to create a balanced control test pool of the same size in which we randomly sampled correct trials from the trials in condition *c* for each neuron *n*. All remaining correct trials of each neuron in *N*_*c*_ (excluding those selected for the control test pool) were used to compute the single-neuron 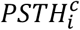 matrix. Here, the superscript *c*, indicates that this is the PSTH of neuron *i* specifically for condition *c*, distinguishing it from the equivalent matrix described for the regular decoder.

For condition *c*, we randomly sampled a single error trial ***e***_*n*_ ∈ ℝ^*T*^ and a single correct trial ***c***_*n*_ ∈ ℝ^*T*^ for each neuron *n* ∈ *N*_*c*_, where *T* is the length of the temporal window. These vectors were concatenated across all neurons to form the error population vector 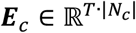 and the correct population vector 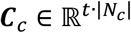.

The data-constraint population matrix 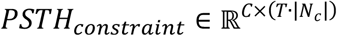 was constructed by concatenating the 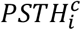 matrices of all neurons in *N*_*c*_, where *C* = 20 is the number of conditions. Each row of *PSTH*_*constraint*_ corresponds to the average population response for one condition.

The Euclidean distances *d*(***E***_*c*_, *PSTH*_*constraint*_[*c*]) and *d*(***C***_*c*_, *PSTH*_*constraint*_[*c*]) were computed for each condition *c*^′^ ∈ {1, ⋯, C}. These distances quantify how closely the sampled population responses (error and correct) match the condition templates in the population matrix. As with the regular decoder, each sampled population vector (***E***_*c*_ or ***C***_*c*_) was classified into the condition *ĉ* with the minimum Euclidean distance:

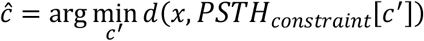

Repeating this process across multiple iterations for each condition *c* resulted in confusion matrices for the control decoder that separately evaluated the classification performance on error and control test groups.

#### Linear Regression Analysis

We applied a leave-one-out linear regression approach to examine the dynamics of SNpr activity along a specific axis.^22^ Below, we detail the methodology, including pre-processing steps, regression modeling, and population-level analysis.

#### Data Pre-Processing

For each iteration and each neuron, we sampled one trial from each condition as a test set. The remaining trials were used to construct a data matrix for each neuron:

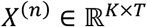

where *X*^(*n*)^ is the data matrix for the *n*-th neuron, *K* is the number of training trials, and *T* is the number of time bins. We used a bin size of 200 ms for this analysis. Each entry *x*_*k,t*_ represents the spike count of the *k*-th trial in the *t*-th time bin. Another matrix was created for the test trials:

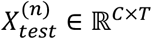

where *C* = 20 represents the number of conditions.

To standardize the data, each entry in *X*^(*n*)^ and 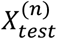 was z-scored using the mean and standard deviation of *X*^(*n*)^:

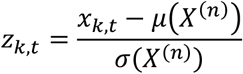

where:

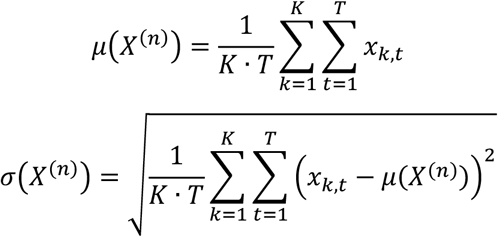

#### Trial Condition Representation

We constructed three standardized vectors describing the trial conditions for the experimental factors:

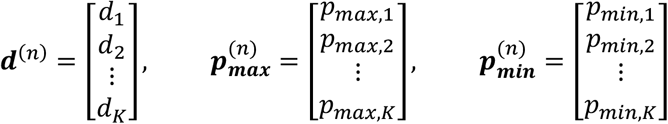

where ***d***^(*n*)^, 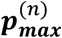 and 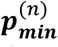 represent the direction, maximal probability, and minimal probability values for all *K* trials of the *n*-th neuron, respectively.

#### Regression Model

For each time bin, we fit a linear regression model to the neuronal activity in the *n*-th neuron:

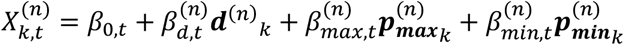

where 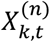 is the spike count for the *k*-th trial in the *t*-th time bin, *β*_0,*t*_ is the intercept, and 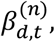,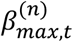 and 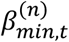 are coefficients for the respective experimental factors.

#### Population Coefficients

For each time bin each in each neuron, we computed the regression coefficients 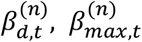 and 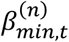, capturing the contributions of direction, maximal probability, and minimal probability. The coefficients of each experimental factor were organized into matrices of size ℝ^*N*×*T*^, where *N* is the number of neurons and *T* is the number of time bins:

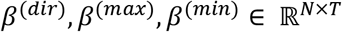

#### Population Activity Projections

We constructed a three-dimensional matrix, denoted as *Test* ∈ ℝ^*C*×*T*×*N*^, which contained the test activity of the entire neuronal population for all conditions, time bins, and neurons. Note that

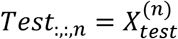

Next, for each condition *c* and time bin *t*, the test activity of all neurons was projected onto the population’s coefficient vector corresponding to direction, 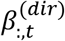, as follows:

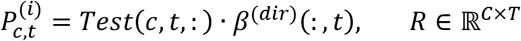

where each entry 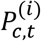 represents the projection of the test activity for condition *c* at time bin *t* during the *i*-th iteration. Importantly, similar results were obtained using a consistent beta across all time points (Figures S5), suggesting that simple linear readouts of SNpr activity can drive behavior.

We repeated this procedure over multiple iterations (*I* = 100), where in each iteration, new test trials 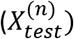 were sampled for each neuron, and the population coefficients (*β*^(*dir*)^, *β*^(*max*)^ and *β*^(*min*)^) were recalculated using the remaining training trials 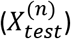.

Finally, we averaged the projection values across all iterations for each condition *c* and time bin *t*:

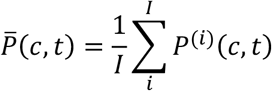

The resulting projection data 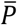 were used to visualize population activity dynamics, with each row corresponding to a condition (*c*) plotted as a separate curve. The horizontal axis represents the midpoint of each time bin (*t*). These plots, presented in Figures 3F and S4, highlight the temporal dynamics of SNpr activity along the decision direction axis.

#### Defining Orthogonal Probability and Direction Axes

To define the probability and direction axes, we first evenly split the trials for each neuron into a test set and a training set. We then fit each neuron’s training set with a linear regression model, as described in the *population activity projection* section.

For the direction axis, we used neural activity from the initial stage of motion (100 to 300 ms after motion onset), since SNpr activity during this time closely matched eye behavior (Figures 1B and 3F). For the probability axis, we used activity from the initial cue stage (100 to 300 ms), since SNpr activity during this period showed the strongest coding of maximal probability (Figure 4B).

After computing these two axes, we applied the Gram-Schmidt procedure to ensure their orthogonality. We then projected the test set activity during the cue and motion epochs onto each of these axes. The probability axis captured the maximal probability of the test set during both the cue and the motion epochs, whereas the direction axis discriminated between different direction conditions in these epochs. Thus, direction and reward information could be easily and independently extracted from SNpr activity. The results of this analysis are presented in Figure S9.

---

**Figure S1:**
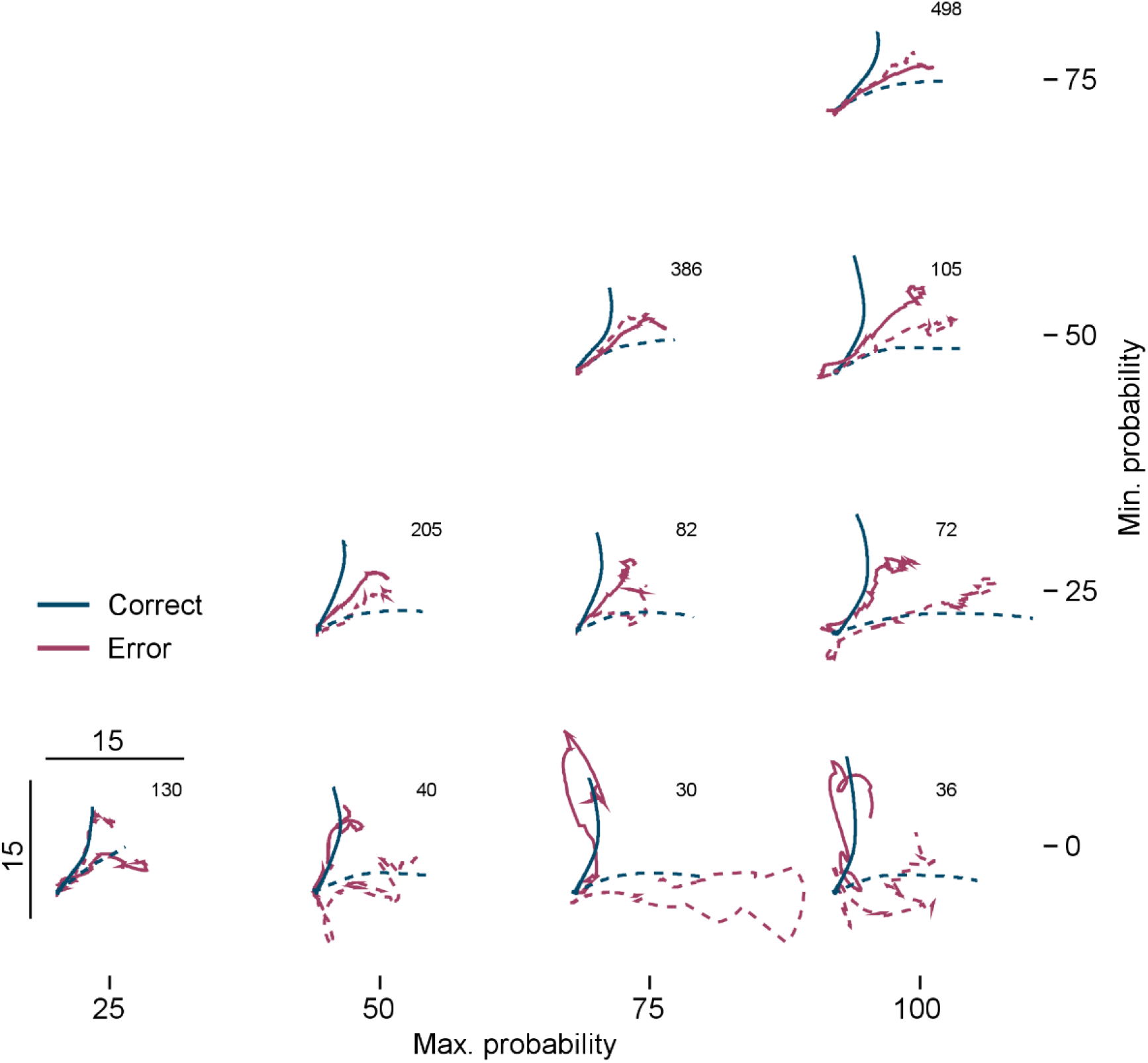
Behavior during error trials. Average eye velocity in the horizontal vs. vertical directions during the first 300 ms after motion onset. Each subplot compares eye velocity for error trials (red) and correct trials (blue) for specific probability pairs indicated by the row and column. Solid and dashed lines represent conditions where the higher-probability target moved vertically and horizontally, respectively. Calibration bars (bottom-left plot) indicate eye velocity of 15°/s. The number at the top of each panel indicates the number of error trials for this reward condition.

**Figure S2:**
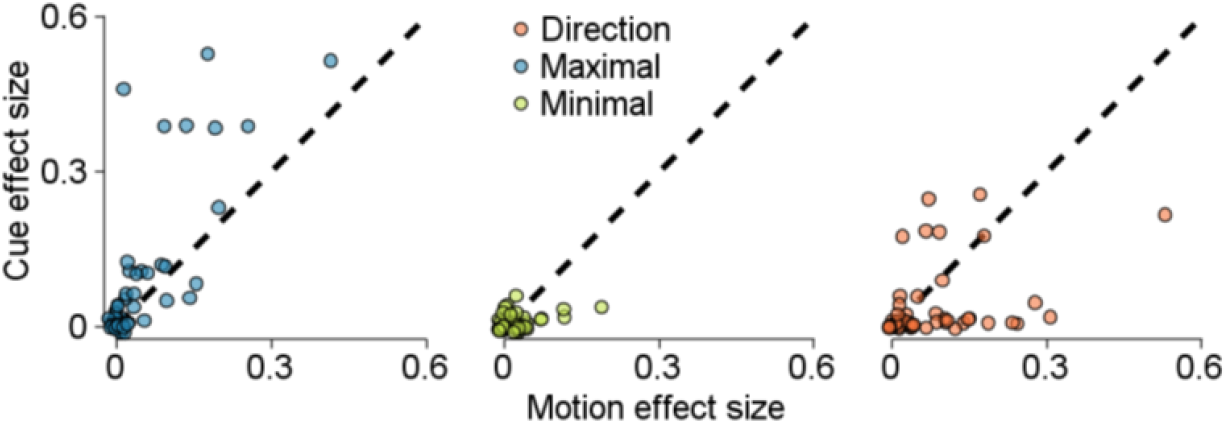
Effect size comparisons during motion onset and cue epochs. Effect size comparison for each experimental factor during the first 400 ms of the cue stage (vertical axis) and motion stage (horizontal axis). Colors indicate different experimental factors.

**Figure S3:**
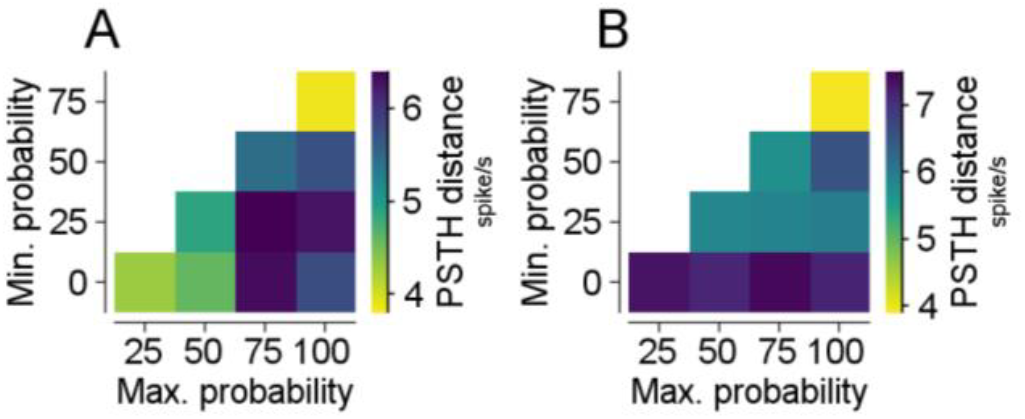
PSTH similarity within task epochs. Same as measured during the cue (**A**) and the initial movement (**B**). Euclidean distance between PSTHs for direction conditions within each reward condition. The maximal and minimal probabilities are plotted on the horizontal and vertical axes. The window for the cue is 0-400 ms from cue onset, and 50-250 ms from target motion onset for movement. Units are spikes per second (see Methods, PSTH distance).

**Figure S4:**
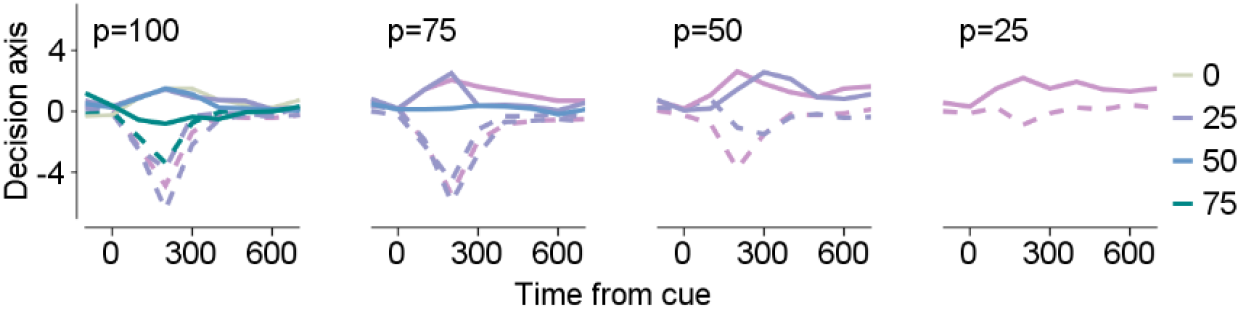
SNpr dynamics on decision axis during cue epoch. Projection of SNpr population activity onto the axis defined by the coefficients of the direction regressor of the single neurons. The horizontal axis represents time from cue onset. The value at the top of each panel represents the maximal probability, while colors indicate the minimal probability. Solid and dashed lines depict conditions where the higher probability target moved vertically and horizontally.

**Figure S5:**
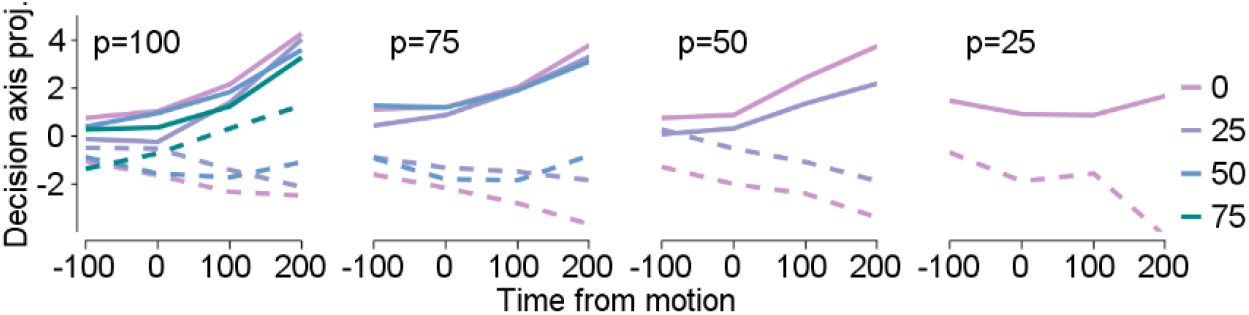
SNpr dynamics on decision axis using linear readout: Projection of SNpr population activity onto an axis similar to Figure 4E but with coefficients fixed across all time points. The horizontal axis represents time from target motion onset, with the value at the top of each panel indicating the maximal probability and colors representing the minimal probability. Solid and dashed lines correspond to conditions where the maximal probability target moved vertically and horizontally, respectively.

**Figure S6:**
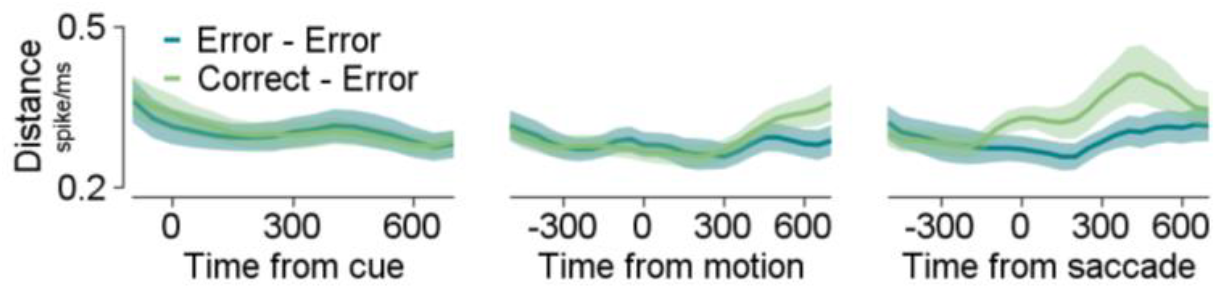
SNpr dynamics in error trials. The Euclidean distance (vertical axis) over time is shown for different trial stages (same as **5C**), comparing the Correct-Error group with the Error-Error group (see Methods) during the cue stage (left), motion stage (middle), and relative to the selection-saccade (right). The Correct-Error and Error-Error groups exhibited similar dynamics during the cue stage (permutation test, p=0.9) but showed differences during the motion stage (permutation test, p=0.01) and the saccade stage (permutation test, p=0.004).

**Figure S7:**
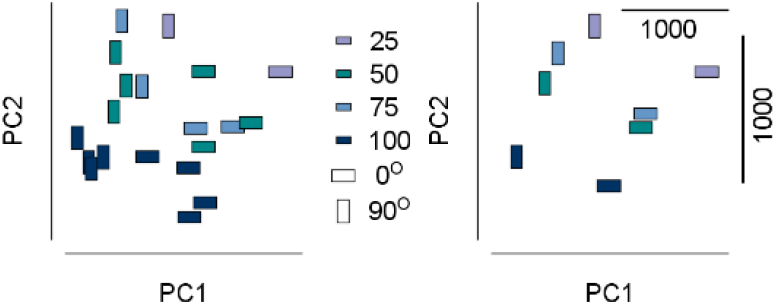
SNpr activity during the pre-commitment stage exhibits an intermediate pattern between cue and movement stages. Similar to Figure 2C,D, this panel shows the PCA projection of SNpr population activity during the pre-commitment stage, calculated from −300 to −100 ms relative to the selection saccade. Left: PCA projection onto the first two principal components across all task conditions. PC1 accounts for 32% of the variability, while PC2 accounts for 15%. Color indicates the maximal probability of each condition, and the rectangle angle reflects the direction condition. Right: Same PCA projection as on the left, with each rectangle’s position indicating the mean position of matched rectangle colors from the left panel.

**Figure S8:**
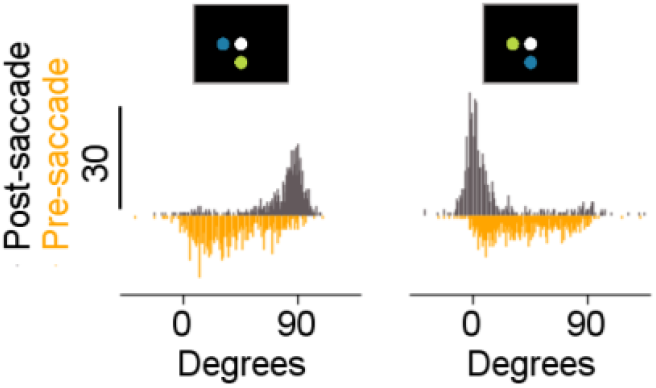
Eye velocity angle during error trials. Histogram of eye velocity angle (degrees) calculated over 200 ms before (yellow) and after (gray) saccade. The inset screen diagram displays the direction condition (horizontal or vertical), with the blue target representing the maximal probability.

**Figure S9:**
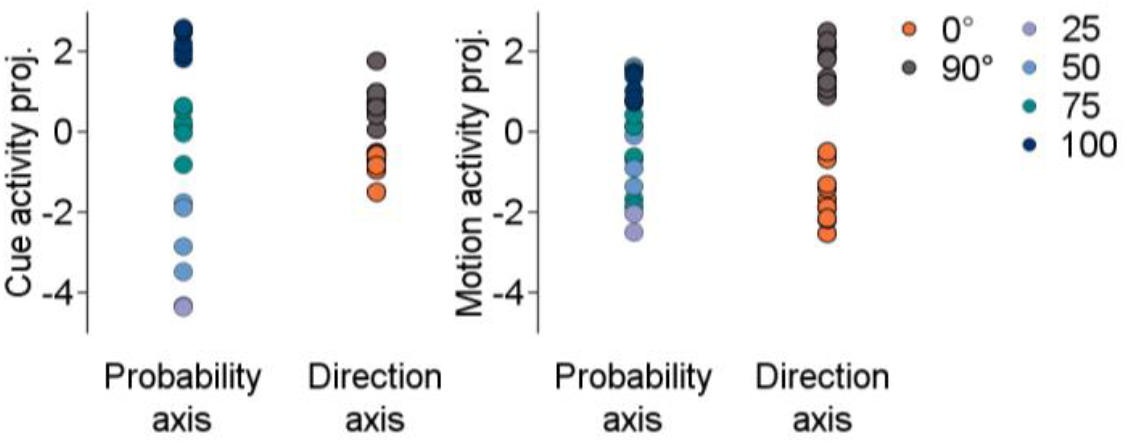
Reward and movement are coded along orthogonal axes. Projection of activity during the cue (left panel) and motion (right panel) stages onto probability and direction axes (see Methods subsection *Defining Orthogonal Probability and Direction Axes*) for all task conditions. Colors indicate maximal probability when projecting onto the probability axis and direction condition when projecting onto the direction axis. The results of this analysis are presented in Figure S9.

